# Context and Phase Dependent Effects of Delta transcranial Alternating Current Stimulation on Dynamic Attending

**DOI:** 10.1101/2021.01.17.427013

**Authors:** Adam Shelp, Giovanna Mioni, Martin Wiener

**Affiliations:** Department of Psychology, George Mason University, Fairfax, VA; Dipartimento di Psicologia Generale - University of Padova – Padova, Italy

**Author notes:** **Corresponding Author: Martin Wiener, PhD**, George Mason University.

## Abstract

Attention requires the allocation of limited resources to properly interpret our environment, making it ultimately unsustainable. Dynamic Attending Theory suggests that, in order to realistically maintain vigilance to our surroundings, attention likely fluctuates between high and low energetic states, such that information can be processed more quickly and accurately during attentional peaks and vice versa. Additionally, prior studies have suggested that the phase of delta oscillations (1-4 Hz) are critically involved in the entrainment of attention. We investigated the physiological and behavioral entrainment of attention and the role that delta phase plays to moderate the benefits of this attending. Participants (N=28) passively listened to a background auditory rhythm and were required to complete a visual discrimination task while undergoing 2 Hz transcranial alternating current stimulation (tACS). The task involved identifying an image, either upright or inverted, presented either on or before the final beat, while receiving delta stimulation that was either aligned or unaligned with image presentation. As expected, reaction times (RTs) were faster for on-beat than off-beat stimuli, and for upright images than inverted. Crucially, tACS phase-aligned with the beat led to faster RTs over out-of-phase stimulation, but only for upright images; remarkably, this pattern was reversed for inverted images presented on-beat, with slower RTs for inverted stimuli during in-phase tACS. These results suggest that the effects of delta tACS are both phase and context dependent, and mediate a potential form of speed-accuracy tradeoff in the allocation of attentional resources during rhythmic entrainment.

## Introduction

Attention is one of our most limited resources, yet is required to complete most daily activities. While we are able to temporarily allocate these reserves to multiple sources, often referred to as “multitasking,” research has shown that this division is ultimately unsustainable (Jefferies & Witt, 2018). As a means of surmounting limited resources, one suggestion is that the brain employs a rhythmic fluctuation of attention over time, such that attention shifts between high and low energetic states. This idea, termed Dynamic Attending Theory (DAT), was originally posited in the late 1970s by Mari Jones, who later suggested that attentional oscillations could be entrained via the introduction of isochronous beats and could stretch and shift based on the statistics of the environment (Jones, 1976; Jones, Boltz, & Kidd, 1982; Jones & Boltz, 1989). More recent work has connected attentional modulation to endogenous neuronal oscillations in alpha and delta bands; moreover, these rhythms can be aligned to exogenous rhythmicity so as to maximize attentional energy at predictable onsets and so potentially enhance perception (Escoffier, et al. 2010; VanRullen, 2016; Nobre & van Ede, 2018 Obleser & Kayser, 2019). However, whether or not internal oscillations do in fact serve to enhance perception or merely redirect it is unknown, necessitating methods for manipulating oscillations to investigate their impact.

Despite increasing interest in DAT, recent work has called into question it’s generalizability. For example, recent work conducted by Kunert and Jongman (2017), which tested participants on a lexical-decision task in which they decided whether a string of letters was a word or non-word, appeared to show that DAT-predicted effects only extend to auditory-motor synchronization rather than selective attention as a whole. Notably, results of another study suggest that the pitch comparison task employed in more recent research conducted by Jones and colleagues (2002) to support DAT was inadequate and did not appropriate demonstrate auditory dynamic attending at all (Bauer, Jaeger, Thorne, Bendixen, & Debener, 2015).

Alternatively, researchers Sanabria, Capizzi, and Correa (2011) used a modified version of the temporal orienting paradigm to test whether attention could be oriented to specific moments by a rhythm. They found that reaction time improved in both long and short intervals, which suggests a highly flexible mechanism behind dynamic attending (Sanabria et al., 2011). These results are further corroborated by the idea that natural stimuli tend to occur in rhythmically consistent patterns (e.g. biological motion or vocal communication) along which attention can be entrained through the concurrence of the excitability phases of oscillations with the timing of event onsets (Schroeder & Lakatos, 2009). Finally, additional research conducted by Triviño and colleagues (2011), and more recently by Morillon and Baillet (2017), quantified this phenomenon by demonstrating that rhythmic attending was localized to the right hemisphere and was accompanied by a shift in delta and beta oscillations.

To properly understand dynamic entrainment, one must be able to definably and quantifiably differentiate between attention and expectation. While most researchers can separately characterize the two by their definitions, however, there is a severe lack of computational and empirical work to explore the mechanisms behind expectation, much of which has been focused on its interaction with vision (Summerfield & Egner, 2009). More recent work has begun to determine the precise neural mechanisms behind improved reaction time in response to more predictable stimuli finding entrainment of low-frequency (1-4Hz) delta oscillations as predictability increases (Stefanics et al., 2010). This was further explained by Rohenkohl and colleagues (2011), who found that enhanced temporal expectations could modulate perceptual processing by altering the contrast sensitivity of visual targets, which was paired with improved reaction times, as well as phase synchronization of delta oscillations. In tandem, these results suggest that expectation is a function of attention, in which the latter is highly focused toward a point in time to create the former, but that there is, indeed, a mechanistic difference between the two.

Research has shown that there may be associations between different frequency bands and time perception across specific contexts (Wiener & Kanai, 2016) despite no direct demonstration in the modulation of transmission efficacy (Fehér, Nakataki, & Morishima, 2017). Following this logic, delta oscillations – which have been shown to arise during many tasks requiring prediction and expectation (Arnal, Doelling, & Poeppel, 2015; Arnal & Giraud, 2012; Lakatos, Karmos, Mehta, Ulbert, & Schroeder, 2008) – should allow researchers to reliably predict and produce changes in attending. This is observed in the results of Henry and Obleser (2012), which demonstrated that perceptual processing of near-threshold stimuli could be affected by oscillatory phase reorganization. Additionally, it has been shown that phase entrainment can enhance sensory processing of visual targets by augmenting their contrast sensitivity (Cravo, Rohenkohl, Wyart, & Nobre, 2013), similar to the findings of Rohenkohl and colleagues (2012).

If delta oscillations are causally involved in shifting dynamic attention, then their manipulation should also alter perceptual functions. One method for influencing endogenous oscillations is transcranial alternating current stimulation (tACS) (Herrmann, Strüber, Helfrich, & Engel, 2016). tACS has been shown to influence perceptual functions and ongoing oscillations (Vosskuhl, et al. 2018) and additionally has been demonstrated in shifting attentional engagement (Hopfinger, Parsons, & Frohlich, 2017). In order to test if delta oscillations are indeed involved in shifting dynamic attention, we stimulated subjects with tACS at 2Hz while performing an image discrimination task embedded in a rhythmic context, in which presented stimuli could occur either at or before an implied beat. Previous work with this paradigm has demonstrated that images presented in-synchrony with the beat are processed faster and accompanied by larger evoked potential (Escoffier, et al. 2010; 2015). We hypothesized that tACS could either enhance or reduce this effect, depending on whether the alternating current was also in-sychrony or not with the implied beat. As described below, our findings only partially confirmed this hypothesis, instead demonstrating that the effect of tACS alignment crucially depended on the difficulty of the discrimination; aligned tACS enhanced easy discrimination but impaired difficult discrimination.

## Method

### Participants

Our sample included 28 participants (13 females, 15 males, mean age: 22.07 years). Participants were right-handed, physically and neurologically healthy adults recruited from the George Mason University campus with ages ranging from 18-35 years old. All subjects had normal or corrected-to-normal vision. Exclusionary criteria included: any previous history of head injury, seizures, migraine, neurological or psychological disorders, and alcohol or drug abuse. Participants also could not be pregnant, nursing, or possibly pregnant. The scalp of each participant was visually inspected immediately preceding the tACS procedure and had any irritation been identified, that subject’s participation would have been postponed or terminated at researcher discretion; no participants were disqualified for this reason. All subjects filled out and signed a consent form prior to initializing the study; the protocol was approved by the George Mason University Institutional Review Board.

### Design

We used a 4-factor within-subjects design with stimulus identity, orientation, timing, and tACS phase as the independent variables. Each factor included two levels: stimulus identity included either face or house stimuli, orientation included upright or inverted images, timing included in-synchrony or out-of-synchrony, and tACS phase included aligned or unaligned. All are described in more detail below.

### Materials

We used a task similar to that of Escoffier, Sheng, and Schirmer (2010). Participants were asked to perform a rhythmically-cued image discrimination task, in which they were required to indicate if a presented image was a house or a face. On a given trial, subjects began by viewing a central fixation cross on a uniform gray background. A rhythmic beat began at the start of each trial, consisting of three snare drum sounds followed by three bass drums (Figure 1c) with an implied beat of 1.3Hz. After the last sound, a silent interval of 1.1s began, during which the visual image stimulus was presented (Figure 1a). Image presentation consisted of either a grayscale face or house stimulus, either upright or inverted (Figure 2a). Subjects were required to respond as quickly but as accurately as possible concerning the category of the image. Each presented measure thus comprised a 3-second interval. At the end of each measure, a second measure began immediately; measures looped in this manner throughout each block. However, image stimuli were only presented in every other measure. A CHERRY MX Red STRAFE Mechanical Gaming Keyboard was used to collect all responses (polling rate: 1000Hz). The ‘F’ and ‘J’ keys were covered by green and yellow stickers respectively and separately assigned to house and face stimuli randomly across subjects with counterbalancing. Subjects completed 24 trials for each condition, across two blocks of 192 trials, for a total of 384 trials. Task design and presentation was programmed using PsychoPy (Peirce, 2007).

**Figure 1.**
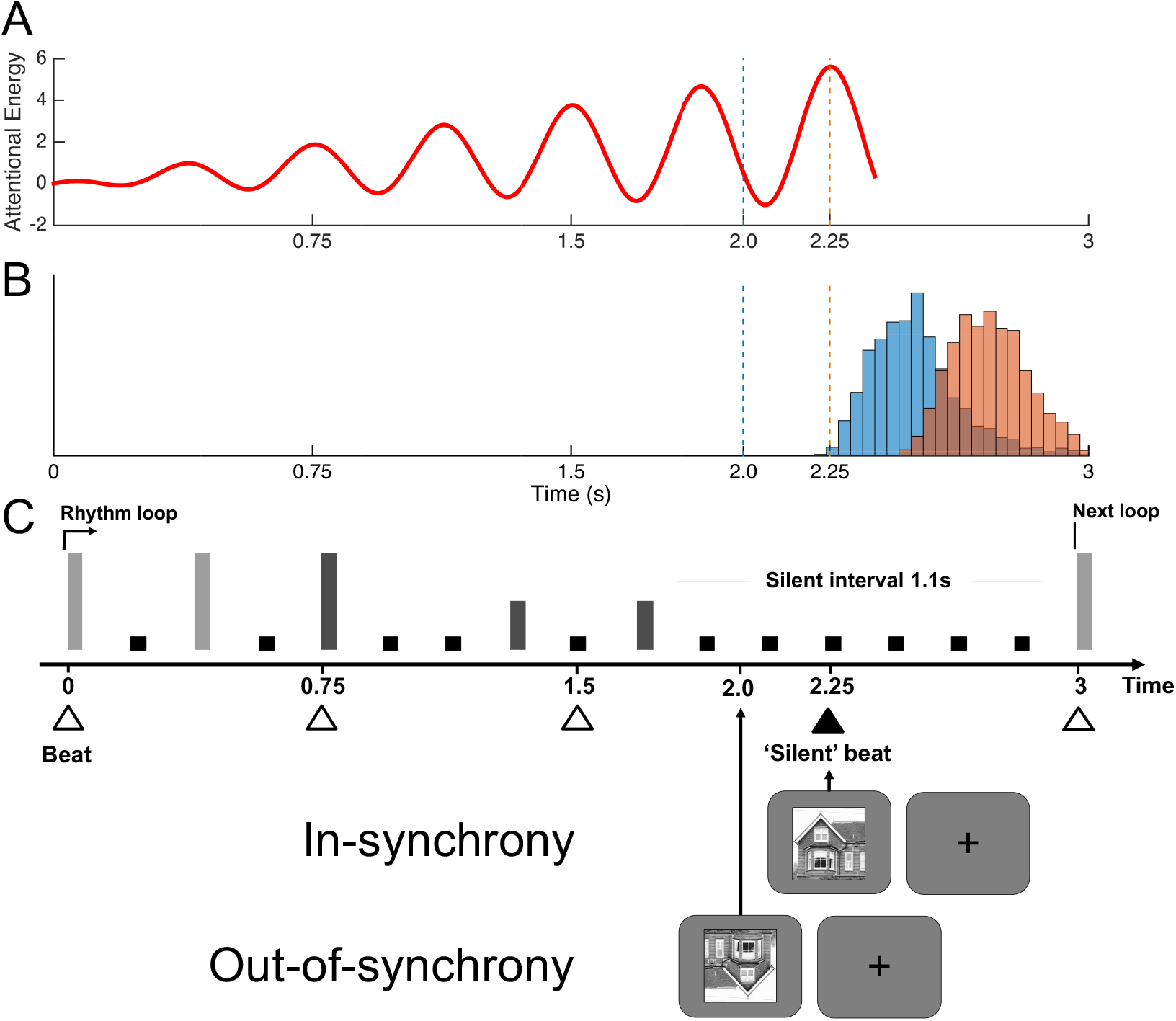
Study design. Subjects were presented with an auditory rhythm loop (adapted from Escoffier, et al. 2010), during which subjects were presented with a four-beat rhythm in which the final beat was silent. An image was presented either in-synchrony or out-of-synchrony with the final beat, which subjects were required to respond to. **A)** Hypothetical change in attentional energy (hypothetical units) in which the rhythm beats drive attention towards the arrival of the fourth “beat” for the expected arrival of the stimulus such that attention is maximal at this point. **B)** Reaction time histograms pooled across all subjects relative to the onsets of in-synchrony stimuli (orange bars and vertical line) and out-of-synchrony stimuli (blue bars and vertical line). **C)** Visual representation of the rhythmic sequence used in the experiment; vertical bars represent sound placement and height represents amplitude. Light gray bars denote snare drum while black bars denote bass drum sounds. Four dots comprise one beat. One measure (four beats, 3,000 ms) and the first beat of the next measure are shown (†=in-synchrony arrival time; *=out-of-synchrony arrival time). Bottom panels indicate visual stimuli presented, in which subjects viewed a blank screen with a fixation cross until the image was presented for 250ms.

**Figure 2.**
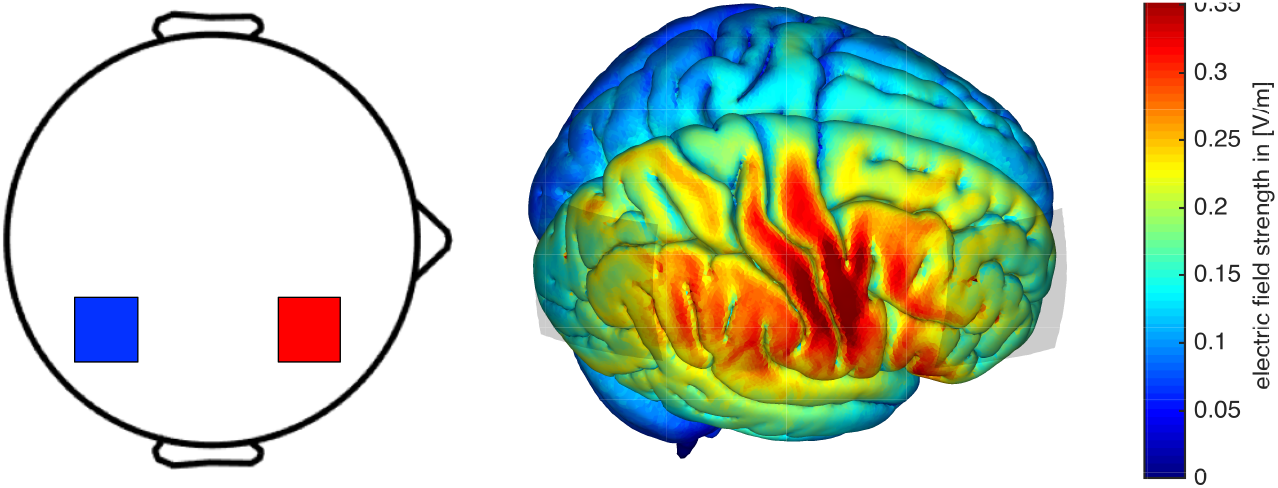
**A)** Stimuli used for the behavioral experiment. Subjects were presented with two exemplars of face or house stimuli, either upright or inverted. **B)** Left: placement for tACS electrodes over F4 (red) and P4 (blue). Right: simulation results from simnibs demonstrating maximal electrical field strength across the right hemisphere.

While completing the task, participants underwent tACS to either entrain delta oscillations along the downbeats or the upbeats of the rhythm. tACS was administered via a battery-powered Neuroconn unit (Neuroconn DC Stimulator+) using two saline-soaked 5×7cm electrodes in sponges at 1.5mA. A 2Hz sine wave was used for all stimulation, and the onset of stimulation was aligned to the first beat of a given measure; however, the phase of the oscillation was shifted by 90° such that an oscillation peak would occur either at 2s or 2.25s, coinciding with the onset of either of the possible images. This shift in stimulation varied between blocks, with subjects performing a single block of aligned or un-aligned stimulation, with the order counterbalanced between subjects. Stimulation terminated at the end of the measure in which an image was presented and was reset at the start of the second measure after; that is, stimuli were presented on every other measure, and tACS only occurred in those measures. Control of the tACS unit was administered via a Labjack U3 interface controlled by Python code within PsychoPy.

To examine the electrical field distribution of the effect we simulated electrode positions using SimNIBS software (version 2.0; http://simnibs.de; Saturnino, et al. 2019) with a current strength of 1.5mA. The normalized electrical field was simulated via a realistic finite element head model. The results of this analysis revealed a broad effect over the right hemisphere between the two electrode sites encompassing fronto-parietal cortex. The right hemisphere was chosen to accord with recent findings that rhythmic modulation and timing are stronger in the right hemisphere (Wiener, et al. 2010; Trivino, et al. 2011; Morillon & Baillet, 2017).

A sensation questionnaire was verbally conducted a minimum of three times throughout the session to ensure participant comfort, with a threshold of 7/10. Any rating of 7 or higher would have resulted in discontinuation from the study, however no participant rated higher than a 6. Finally, a post-task questionnaire was provided at the end to collect participant demographics as well as any significant mood alterations by asking if their mood felt any different post-stimulation. Following drastic changes, the researcher would have monitored the participant and reevaluated every 15 minutes until their mood had stabilized, however no participant reported drastic mood changes.

## Procedure

Participants were seated in front of a computer monitor (Dell Gaming 27”, 100Hz refresh rate) before completing an informed consent form. Once the form had been collected, the researcher attached electrodes over areas F4 and P4 to stimulate the across the right hemisphere (Figure 2b). After the electrodes were placed, but before the unit had been activated, the researcher verbally conducted a sensation questionnaire to form a baseline against which future answers could be compared. Once scored, a brief exposure session was run to ensure that participants were prepared for any and all sensations or side effects that might result from stimulation. This exposure lasted 17 seconds total, with a 1-second fade-in, 15 seconds of full stimulation, and a 1-second fade-out. The sensation questionnaire was conducted immediately following completion of this exposure session to ensure that participants were comfortable enough to undergo full stimulation. Before the task began, the sensation questionnaire was verbally conducted once more to ensure participant comfort, after which, the task – which took approximately 15 minutes – began.

Upon completion of the first iteration of the task, participants were allowed to take a short break to recuperate before continuing to the second iteration. This was identical to the first outside of the stimulation phase, which was changed based on the initial condition (in-phase or out-of-phase, counterbalanced between subjects). Afterwards, the participant was asked to complete the post-task questionnaire before departing the study.

### Analysis

Behavioral data for subjects were analyzed by first log-transforming the reaction time (RT) values, thus allowing them to conform to a normal distribution (Baayen & Milin, 2010). Once transformed, RT data were analyzed via a linear mixed effects model (LME), in which RT values were regressed against four fixed effects: Stimulus (Face,House), Orientation (upright, inverted), Timing (Early, On-Time), and tACS Phase (Aligned, Unaligned); Subject was included as a random effect. Only correct trials were used for this analysis. Model fitting was accomplished via Restricted Maximum Likelihood Estimation. Significance of individual effects was assessed by examining 95% confidence intervals for each estimate, as well as Satterthwaite corrected p-values (alpha=0.05). Follow-up tests of interactions were conducted using paired t-tests. An additional analysis was conducted on accuracy data by calculating the average percent correct for each condition. These responses were analyzed via a repeated measures ANOVA with stimulus timing and tACS phase as within-subject factors.

## Results

All subjects were able to perform the task well without any major difficulties. A repeated-measures ANOVA of task accuracy (percent correct) found no difference across stimulus timing or phase conditions (Average percent correct = 92% ± 8% (SD); all *p*>0.05) (Figure 3c).

**Figure 3.**
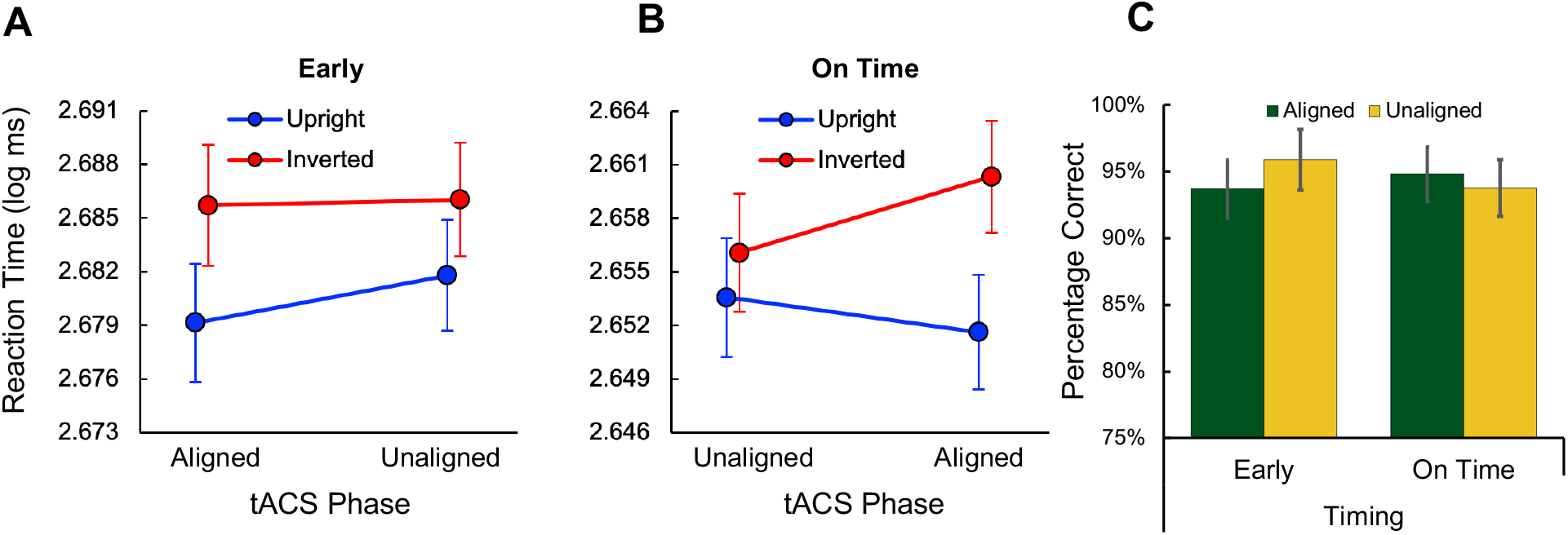
Behavioral results of stimulation. **A)** When the image was presented early (out-of-synchrony with rhythm) reaction time values were significantly slower for inverted stimuli compared to upright, regardless of whether tACS was applied aligned or unaligned with stimulus onset. **B)** When the image was presented on time (in-synchrony with rhythm) reaction times diverged depending on tACS phase and image presentation, with faster reaction times when upright stimuli were aligned with tACS, but slower reaction times when inverted stimuli were aligned with tACS. **C)** Accuracy of responses across timing, stimulus and tACS conditions; no effects were observed across any condition on the accuracy of responses. Error bars indicate standard error.

The results of the LMM analysis revealed a significant effect of stimulus timing characterized by faster RTs when the stimulus occurred on time [β = −0.096, 95% CI: −0.13 – 0.06, *p*<0.001], thus replicating the previously reported effect (Escoffier, et al. 2010). However, no main effect was found of either orientation [β = 0.102, 95% CI: −0.011 – 0.216, *p*=0.08] or stimulus identity (face or house) [β = 0.0194, 95% CI: −0.093 – 0.132, *p*=0.73612]. Additionally, we found no main effect of tACS phase on RT [β = 0.0006, 95% CI: −0.0005 – 0.0019, *p*=0.30224]. However, we did observe a significant interaction between orientation and phase [β = −0.0019, 95% CI: −0.0036 – 0.00011; *p*=0.0372] as well as a three-way interaction between orientation, timing, and phase [β = 0.0009, 95% CI: 0.00008 – 0.0017158; *p*=0.0316]. No other interaction effects were significant (all *p*>0.05). Examination of mean RT between timing and orientation conditions revealed a differential effect of tACS phase on RT. Specifically, we found that when the stimulus was presented early, upright images were generally faster than inverted images, but there was no effect of tACS at either phase alignment (Figure 3a). In contrast, when the stimulus was presented on time, upright images demonstrated the anticipated effect of improved reaction times with aligned tACS, however, when the stimulus was inverted, a decreased reaction time was observed for aligned tACS [paired *t*-test of upright vs inverted: *t*(27)=3.243, *p*=0.003, Cohen’s *D* = 0.612] (Figure 3b).

## Discussion

The present study tested two primary hypotheses. The first was that individuals who are stimulated at a delta frequency that aligns with the downbeats of the presented rhythm will demonstrate faster reaction times associated with images presented in-synchrony than those presented out-of-synchrony. The second was that individuals who are stimulated at a delta frequency that aligns with upbeats of the presented rhythm will demonstrate faster reaction times associated with images presented out-of-synchrony than those presented in-synchrony. Our results only partially supported the latter hypothesis, but additionally provided some novel findings.

As expected, we replicated the findings of Escoffier and colleagues (2010), which show that musical rhythms synchronize and enhance visual stimulus processing, regardless of stimulus type, and further suggests that DAT can effectively entrain along an implied isochronous rhythm. Specifically, subjects were faster in responding to visual stimuli that occurred at the implied fourth beat in the measure than when the stimulus occurred 250ms earlier. Our prediction was that delta-tACS would enhance this effect at the implied beat when the current peak was aligned with visual stimulus onset, but would also shift this improvement to the earlier onset when the peak was shifted into alignment with this earlier moment. Notably, subjects were not given any instruction regarding the onset of visual stimuli or the musical rhythm, only being told they were required to respond to the images as fast as possible and that there would be a background rhythm. Instead, we found no effect of tACS at the earlier timepoint. One possible reason for this is that the influence of the auditory rhythm for entrainment had a greater impact than tACS. Indeed, tACS can only weakly influence ongoing oscillations (Vöröslakos, et al. 2018), and there is controversy regarding the route with which tACS influences neuronal processes, with some findings suggesting a route via cranial nerves (Asamoah, et al. 2019). In contrast, auditory entrainment is a robust phenomenon that can have a strong influence on oscillatory power (Lakatos, et al. 2008; Lakatos, et al. 2019). Indeed, the impact of stimulus timing on RT was an order of magnitude larger than the tACS effect that aligned with the rhythm, further suggesting that the capability of tACS for shifting exogenously-entrained rhythms is low.

Contrary to the expected results, we found that the impact of phase-aligned tACS with the implied beat had a differential impact, depending on the difficulty of the discrimination. We observed that subjects tended to be slower for inverted as opposed to upright stimuli, although this effect was not significant. Yet, phase-aligned tACS with the implied beat shifted the RT for each in opposite directions, with RT slowing down for inverted images but speeding up for upright ones. This finding suggests that the effect of phase-aligned tACS is context dependent, such that its influences crucially depends on the stimuli being processed at the entrained moment.

Why might phase-aligned tACS shift response time in opposite directions for upright and inverted stimuli? According to a general assumption of both DAT (Henry & Hermann, 2014) and the effects of tACS (Vosskuhl, et al. 2018), phase-aligned stimulation should enhance attention at the implied beat and so lead to faster responses for both inverted and upright stimuli. However, this also assumes that 1) upright and inverted stimuli are processed equivalently, and 2) the effects of tACS are constant across conditions, neither of which may be correct. Indeed, previous work has shown that inverting an image significantly alters categorization independent of detection (Mack, et al. 2008). For tACS, more recent findings have also demonstrated state-dependency in its effects (Alagapan, et al. 2016; Lefebvre, et al. 2017), suggesting that the behavioral state of the subject can alter the impact of tACS. With respect to the present findings, one possible interpretation is that tACS-induced changes in RT are mediated by a stochastic resonance type mechanism (Schwarzkopf, et al. 2011). In this framework, one can construe inverted stimuli as “noisy” in that they require additional resources to categorize. Similarly, low-frequency tACS may in fact act by boosting delta oscillations, which themselves act to coordinate larger “bursts” in higher-frequency bands (Arnal, et al. 2015), which in itself may be a kind of noise. In combining these two together, the presence of an inverted, noisy, stimulus with increased energy from tACS may in fact boost the noise, whereas the presence of an upright, clearer, stimulus is enhanced. This finding would further imply an inverted-U shape effect of tACS with task difficulty, such that there exists an optimal level of stimulation for enhancing perception. Similar findings have been demonstrated for transcranial random noise stimulation (tRNS), where an appropriate intensity is necessary to enhance the perception of subthreshold visual stimuli (van der Groen & Wenderoth, 2016).

While the above findings suggest caveats to stimulation and a mechanism that can explain our findings, we note limitations to the present results. First, we note that, given the weak effect of tACS, it is unclear if the observed effects were truly due to an enhancement of entrained oscillations or a different, non-specific effect. Second, an alternative reason for the discrepant findings could be due to our choice of electrode montage. To this latter alternative, we originally decided to stimulate the right hemisphere due to previous work suggesting a right-hemispheric advantage for rhythmic processing (Morillon & Baillet, 2017). However, another possible stimulation site could have been over the visual cortex itself, representing the endpoint of dynamic attending for enhancing image processing (Escoffier, et al. 2015). As such, we cannot rule out the possibility that stimulating over this region would have in fact led to the predicted effects, or an enhanced version of the observed findings. Future work will be necessary test these differences.

Altogether, our findings confirm that rhythmic entrainment can enhance the processing of visual images, but also suggest that this effect is far larger than the influence of tACS. Further, they suggest that tACS may have state dependent effects depending on the difficulty of the task employed, such that stimulation could either enhance or diminish processing of visual stimuli. These findings present context and phase-specific effects for brain stimulation in the context of environmental rhythms.

## Acknowledgements

The authors declare no conflicts of interest. Data from this study are available upon request from the corresponding author.

